# Intraflagellar transport-20 mediates the ciliary membrane trafficking of channelrhodopsin in *Chlamydomonas reinhardtii*

**DOI:** 10.1101/2025.11.29.691277

**Authors:** Alka Kumari, Shilpa Mohanty, Snigdha Samanta, Suneel Kateriya

## Abstract

The primary cilium is a microtubule-based organelle essential for cellular signaling, whose assembly depends on intraflagellar transport (IFT). IFT20, a unique IFT-B component, localizes to both Golgi and cilium/flagellum, and plays a role in ciliary membrane protein trafficking. Here, we analyzed IFT20-mediated ciliary membrane trafficking of channelrhodopsin-1 (ChR1) in *Chlamydomonas reinhardtii*. IFT20 and ChR1 was found to be co-localized throughout the cilia of *C. reinhardtii* wild-type strain. In *bbs1* mutants, IFT20 exhibited distribution along the flagellar length and basal body region. In contrast, ChR1 was found to be accumulating in the distal region of the cilia. Further, protein interaction network analysis reveals that IFT20 serves as a central adaptor, interfacing ancillary ciliary trafficking components, including Arf, the IFT complex, and BBSome subunits. The Arf co-localized with IFT20 in the cilia of the wild-type strain, suggesting a potential interaction. The fluorescence spectroscopic analysis shows that upon GTP binding, IFT20 undergoes concentration-dependent fluorescence quenching. From the far-UV CD spectroscopic analysis, it was observed that recombinant IFT20 has a predominantly helical structure, which does not change upon GTP binding. These findings extend the mechanistic understanding of ciliary membrane protein delivery in *C. reinhardtii* and reveal a conserved role of IFT20 in photoreceptor trafficking.

## 1. Introduction

Intraflagellar transport (IFT) is an efficient coordinated process that drives the bidirectional trafficking of protein complexes in the ciliary axoneme. It comprises of two complexes, IFT-A and IFT-B. The IFT-B complex facilitates anterograde movement of cargo proteins toward the ciliary tip through the activity of the kinesin-2 motor. The IFT-A complex facilitates retrograde movement of IFT particles and associated cargo via dynein-2 motor protein (Pazour et al., 2002, Taschner et al., 2016, Morthorst et al., 2018, Wang et al., 2018).

The primary cilium is a non-motile, microtubule-based projection that functions as a sensory antenna for extracellular signals. Its structure and function rely on precisely regulated trafficking pathways that direct membrane proteins and signaling molecules from the cytoplasm and Golgi to the cilia (Satir et al., 2010, Wingfield et al., 2018). The cargo delivery is governed by specific targeting motifs, and by the coordinated vesicular transport orchestrated through the IFT machinery and ancillary proteins (Rab GTPases, Arfs) (Deretic et al., 2005, Leroux 2007, Ward et al., 2011, Hanke-Gogokhia et al., 2016). The breakdown of any of these processes leads to a disruption of the core signaling pathways leading to a diverse group of developmental and sensory disorders known as ciliopathies (Fliegauf et al., 2007, Reiter and Leroux 2017, Mill et al., 2023). Central to ciliary assembly and homeostasis is the IFT system, a multi-subunit transport machinery composed of two major complexes (IFT-A and IFT-B) (Lechtreck 2015, Jiang et al., 2023). The IFT proteins are localized to the basal body and cilium. However, IFT20 is also found to be localized in the Golgi apparatus in addition to the basal body and the cilia (Follit et al., 2008). This dual localization enables IFT20 to serve as a critical link between post-Golgi vesicular trafficking and the targeting of cargo proteins to the cilium.

This process depends on coordinated interactions with members of the Arf and Rab families of small GTPases (Deretic 2013). In particular, Arf4 contributes to the cargo sorting at the Golgi by recognizing ciliary targeting signals and promoting the formation of transport vesicles carrying ciliary proteins (Deretic et al., 2021). The dysregulation of the Arf and Rab proteins, is detrimental to the functioning of the IFT20 adaptor, thereby affecting the integrity of the IFT complex. This leads to flawed ciliary assembly or dispositioned signaling receptors, which in turn contributes to sensory impairments and ciliopathies. Apart from the IFT machinery, BBSome, an octameric protein complex acts as a critical adaptor for ciliary membrane trafficking. Mutations in BBSome components impair trafficking efficiency, causing IFT proteins and membrane receptors to accumulate abnormally at the ciliary tip, as observed in Bardet-Biedl syndrome. The BBSome comprises multiple subunits with specialized functions. The BBSome is comprised of several essential subunits, such as BBS1, BBS2, BBS7, and BBS9, which create the structural basis of the complex and engage in interactions with cargo proteins associated with the membrane. In contrast, additional components such as BBS4, BBS5, BBS8, and BBS18 contribute to the proper localization of the BBSome at the basal body and support its association with cellular membranes (Klink et al., 2017). The structural characterization has shown that the IFT20-IFT38 module within the IFT-B complex interacts with BBS1, BBS2, and BBS9, creating a functional docking interface either at the base of the cilia or along IFT trains (Nakayama and Katoh 2018). This interface facilitates the regulated transfer of GPCRs and other signaling cargo between the IFT machinery and the BBSome, ensuring precise loading and unloading at the ciliary base and tip (Nozaki et al., 2018). This cargo selection trafficking is found to be conserved across eukaryotes, where the single-celled alga *Chlamydomonas reinhardtii*, shows comparable trafficking processes for the localization of bacteriorhodopsin-type rhodopsins within the flagellar membrane. This plays a crucial role in the light-driven phototaxis (Awasthi et al., 2016). Nevertheless, the role of IFT20 in the trafficking of channelrhodopsins remains poorly understood, representing a significant gap.

To address this gap, we investigated the role of IFT20 in ciliary trafficking of channelrhodopsin-1 in *C. reinhardtii*. Previous studies suggest that channelrhodopsin-1 (ChR1) relocates between the eyespot and flagella in a light-dependent manner, whereas channelrhodopsin-2 (ChR2) is exclusively localized to the eyespot and basal body under similar conditions (Awasthi et al., 2016). Building on this framework, we performed a comprehensive bioinformatic, biochemical, and cell-biological assessment, to analyse the role of IFT20 in the ciliary trafficking of membrane proteins. In this study, we focus solely on the IFT20-mediated ciliary distribution of channelrhodopsins (ChR1). Our study shows the co-localization of the IFT20 and ChR1 throughout the *C. reinhardtii* wild-type strain’s cilia. In *bbs1* mutants, IFT20 exhibited distribution along the flagellar length and basal body region. In contrast, ChR1 was found to be accumulating in the distal region of the cilia. Further, protein interaction network analysis reveals that IFT20 serves as a central adaptor, interfacing ancillary ciliary trafficking components, including Arf, the IFT complex, and BBSome subunits. Further, we also report the potential co-localization of IFT20 with the small GTPase, Arf in the cilia, suggesting a potential interaction.

## 2. Materials and methods

### 2.1 Bioinformatic analysis of IFT20 homologs

The IFT20 protein sequences were acquired from the NCBI (https://www.ncbi.nlm.nih.gov/) and JGI Genome Portal (https://jgi.doe.gov/) by performing pBLAST searches against the non-redundant database. Domain architecture was analyzed using the CDART tool (Geer et al., 2002). The multiple protein sequence homology analysis was performed using the ClustalW (https://www.ebi.ac.uk/jdispatcher/msa/clustalo) program and visualized by BioEdit (http://www.mbio.ncsu.edu/bioedit/bioedit.html). Conserved sequence motifs within IFT20 homologs were detected using the Multiple Expectation Maximization for Motif Elicitation (MEME) Suite (MEME version 5.5.8) (Bailey et al., 2009). Motifs showing strong statistical support (E-value: 1 × 10□□) were selected for downstream analyses, and their significance was determined based on the associated E-values. An analysis of the secondary structure was performed using the CFSSP (Chou and Fasman Secondary Structure Prediction Server) online tool, available via the ExPASy server. Phylogenetic analysis of IFT20 proteins was performed in MEGA11 using 1000 bootstrap replicates (Tamura et al., 2021). Pairwise distances were calculated with the JTT model, and initial trees were generated using the Neighbor-Joining and BioNJ methods. These preliminary topologies were then evaluated to identify the tree with the highest log-likelihood score.

### 2.2 Protein-protein interaction network prediction

IFTs, BBSomes, and ancillary protein sequences were obtained from JGI Genome Portal and NCBI database and subjected to protein-protein interaction (PPI) analysis using STRING v11, based on multisource evidence (Szklarczyk et al., 2019). Interaction networks were subsequently visualized using Cytoscape v3.10.3 (Shannon et al., 2003).

### 2.3 Structural modeling of IFT20 homologs

The tertiary structures of IFT20 homologs were predicted using the AlphaFold Server (https://alphafoldserver.com/welcome). The predicted template modeling (ipTM) score of less than 0.6 from AlphaFold-Multimer suggests random predictions, while a score between 0.6 and 0.8 leads to a performance that exceeds random chance, with improved (Zhu et al., 2023, Homma et al., 2024). The interaction models that were predicted were visualized using PyMOL version 3.1. Similarly, the AlphaFold structure of IFT20-GTP-Mg^2+^ was predicted.

### 2.4 Heterologous expression and affinity purification of recombinant CrIFT20 proteins

The synthesized CrIFT20 coding sequence was sub-cloned into the pET21a vector and expressed in *E. coli* BL21 (λDE3) cells. An overnight-grown culture was inoculated into 500 mL Terrific Broth medium (TB) containing 100 µg/mL ampicillin, and incubation was carried out at 37□ °C with shaking at 200 rpm till the OD□ □ □ 0.6-0.8 was attained. After chilling on ice for 45 minutes, culture induction was carried out with 0.3 mM IPTG and grown at 16 °C for 48 h. Cells were harvested (5000 rpm, 10 min, 4 °C), and resuspended in 50 mM Tris-HCl buffer pH 8 buffer containing lysozyme (0.5 mg/mL) and PMSF (0.2 mM). The cells were lysed by sonication (50 % amplitude, 10 s on/off, 10 min). The lysates were subjected to centrifugation (13,000 rpm, 60 min, 4 °C), filtered (0.45 µm), and the resulting supernatant was loaded onto a pre-equilibrated Ni-NTA IMAC column in 50 mM Tris-HCl (pH 8), 300 mM NaCl, and 20 mM imidazole. After binding and washing, His-tagged CrIFT20 was eluted with 250 mM imidazole. Protein sample’s separation was undertaken by electrophoresis on a 12 % SDS-PAGE gel and then electroblotted onto a nitrocellulose membrane at 4□ °C (350 mA, 90 min). Blocking was done with 5 % skimmed milk in PBST and incubated with anti-His primary antibodies (1:5000), followed by HRP-conjugated secondary antibodies (1:5000). Blot was developed by enhanced chemiluminescence (ECL) using solution A (100 mM Tris-Cl, pH 8.5, luminol and coumaric acid) and solution B (100 mM Tris-Cl pH 8.5 and hydrogen peroxide), and protein bands were captured on photographic films in the dark.

The recombinant IFT20 was used for its biophysical characterization, in the absence and presence of increasing concentrations of GTP (Guanosine triphosphate). The GTP binding assay was selected to analyse the probable binding and conformational change of IFT20 with GTP. In this study, we analyse the interaction of IFT20 with Arf (small GTPase). Hence to study, the possible interaction of GTP with IFT20, we performed GTP-binding assays as described in the next section.

### 2.5 Fluorescence spectroscopy of recombinant CrIFT20

Intrinsic tyrosine fluorescence measurements of recombinant CrIFT20 (dialyzed in 20 mM Tris-HCl pH 8) were performed using a Varian Cary Eclipse fluorescence spectrophotometer (Agilent Technologies). Excitation of samples was done at 275 nm (5 nm excitation slit width), and emission spectra were recorded from 290-500 nm (5 nm emission slit width) at room temperature. GTP (Sigma-Aldrich) was diluted in the reaction buffer (20 mM Tris-HCl pH 8, 3 mM MgCl_2_). Buffer blank fluorescence was subtracted from all spectra, and data were analyzed by standard methods.

### 2.6 Circular dichroism spectroscopy of recombinant CrIFT20

Purified recombinant CrIFT20 was analyzed by far-UV circular dichroism (190-260 nm), using 20 mM Tris-HCl, pH 8 and 3 mM MgCl_2_ as the buffer control at room temperature. CD spectra were monitored following the addition of increasing GTP concentrations. Measurements were performed on a Jasco J-815 spectrometer (Jasco, Tokyo, Japan) containing a 250 W xenon lamp, using a 1 mm path-length cuvette, a scanning speed of 50 nm.min^-1^, and a bandwidth of 0.5 nm.

### *2*.*7 Chlamydomonas* culture, protein extraction, and immunoblotting

*C. reinhardtii* wild-type (CC-125) strain used in this study was obtained from the Chlamydomonas Resource Center (https://www.chlamycollection.org/). Wild-type strain was cultured in standard Tris-acetate-phosphate (TAP) medium. Culture was grown at 22 °C under light/dark cycle of 14 h of light followed by 10 h of dark, at 110 rpm (Catalan et al., 2025). Harvesting was done at 5000 rpm for 10 min, followed by resuspension in 1 X PBS with 0.1 % protease inhibitor cocktail (PIC). Lysis was carried out via sonication at 30 % amplitude with 8 s on/off cycles. Post centrifuging the cells at 13,000 rpm for 15 min, the clarified lysate was collected, mixed with Laemmli buffer, and heated at 65□°C for 30 min. Protein separation was done using SDS-PAGE, and protein bands were transferred to the nitrocellulose (NC) membrane as per the laboratory protocol. Primary antibody diluted at a ratio of 1:1000 in 1 X PBS was used and incubation was carried out for 1 h. Post PBST wash, incubation was carried out using horse radish peroxidase (HRP) conjugated secondary antibody (1:1000) (Sigma Aldrich, St. Louis, Missouri, United States) for 1 h. Blot was developed by enhanced chemiluminescence (ECL) method. The anti-ChR1-Ct and anti-IFT20 antibodies were generated in the laboratory ((Awasthi et al., 2016, Sushmita et al., 2025).

### 2.8 Immunolocalization of CrIFT20 and ChR1

Coverslips were washed and coating was done with 0.1 mg/mL poly-L-lysine (Sigma-Aldrich). Seeding of the cells was done on the coated coverslips and incubation was carried out at room temperature for 15 min. The cells were fixed with a freshly prepared 3.7 % paraformaldehyde in PBS for ∼ 7 min. Following fixation, the coverslips were transferred to chilled absolute ethanol at - 20□ °C for 10 min with occasional gentle agitation. Post fixation and permeabilization, rehydration was done in PBS containing 250 mM NaCl, followed by washing with PBS comprising 0.5 % Triton X-100 (PBST). Incubation was done in blocking buffer (5 % BSA prepared in 1 X PBST) for 1□h at room temperature. The permeabilized cells were incubated overnight at 4□°C with freshly diluted primary antibodies, CrIFT20 (1:250) and ChR1-Ct (1:250). After washing with PBST and PBS, incubation was done at 37□°C for 1 h with Alexa Fluor-conjugated secondary antibodies (1:1000) (anti-goat Alexa-488 and anti-rabbit Alexa-546; Invitrogen, USA). Following additional washes with PBST and PBS, the coverslips were mounted onto glass slides using SlowFade® Gold antifade reagent (Molecular Probes, Invitrogen, USA) in an inverted orientation for imaging (Lechtreck 2013). Samples were imaged using STELLARIS (Leica) confocal microscope at SATHII Foundation, IIT Delhi. The in-house anti-Arf antibody was generated in the laboratory (Ranjan et al., 2014).

### 2.9 Co-immunoprecipitation

°

*C. reinhardtii* cells were grown under a 14 h light/10 hr dark cycle and collected by centrifugation (5000 rpm, 10 min). Cell pellets were resuspended in 50 mM Tris-HCl, pH 8, buffer containing 1 % NP-40, 1 mM PMSF, 1 mM sodium orthovanadate, and 0.1 % SDS, along with a cocktail of protease inhibitors (Sigma-Aldrich). The lysates were sonicated and precleared by incubation with preimmune serum and a protein A/G bead slurry for 2 h at 4□°C with gentle rotation. The separation of the beads from the lysate was carried out by centrifugation at 13,000 rpm at 4 °C for 10 min. For immunoprecipitation, 1 mL of the pre-cleared lysate was incubated overnight at 4□°C with rotation using pre-immune serum (negative control) and the specific primary antibody. The lysate was then incubated with Protein A-Sepharose beads slurry for 2 h at 4 ° C. Immune complexes were harvested at low speed, washed three times with 1 X PBS, and elution was done with 0.1 M glycine (pH 2.8). The eluted samples were mixed with 4X Laemmli sample buffer and heated at 65□ °C for 30 min. The contents were separated by SDS-PAGE, and analyzed by immunoblotting with the indicated antibodies using the procedure described above.

## 3. Results

### 3.1 Phylogenetic and sequence homology analysis reveal a conserved functional core of IFT20 in ciliary trafficking

Analyses of comparative sequence homology and secondary structures of IFT20 homologs demonstrated that the coiled-coil domain at the C-terminus is conserved. Conversely, the N-terminal region shows significant sequence diversity and is composed of β-strands connected by a flexible loop. Motif analysis additionally revealed three distinct, conserved sequence elements amongst homologs, which were identified using expectation-maximization algorithms that evaluate motif enrichment and position-specific conservation **(Fig. 1A)**. These motifs delineate essential interaction sites necessary for engaging with intraflagellar transport components or adaptor proteins related to ciliary trafficking (Boukhalfa et al., 2021). Importantly, these conserved residues are primarily located within predicted α-helical segments. Structural conservation was assessed by superimposing IFT20 homologs with *C. reinhardtii* IFT20 (CrIFT20) serving as the reference **(Fig. 1B)**. The structure validation was done via AlphaFold predicted template modeling score **(Supplementary Table S1)**. Oomycete IFT20 proteins are organized into a distinct clade (bootstrap 100), indicating the adaptation of the IFT pathway within this lineage. In contrast, metazoan homologs form a separate, more basal branch, which aligns with an early evolutionary divergence of the ciliary transport system in animals. Among all the groups, chlorophyta displays the greatest retention of the reference (CrIFT20) IFT20 structure **(Fig. 1C)**, which aligns with the preserved molecular characteristics necessary for ciliary transport in *Chlamydomonas*. Collectively, the combined sequence alignment and phylogenetic analysis reveal that IFT20 maintains a highly conserved core that is crucial for its fundamental role in IFT-B assembly, whereas the surrounding regions show lineage-specific variability that might facilitate specialized regulatory roles. The close relationship of *Chlamydomonas* with other chlorophytes further supports the application of *C. reinhardtii* as an effective model organism for ciliary trafficking studies.

**Fig. 1.**
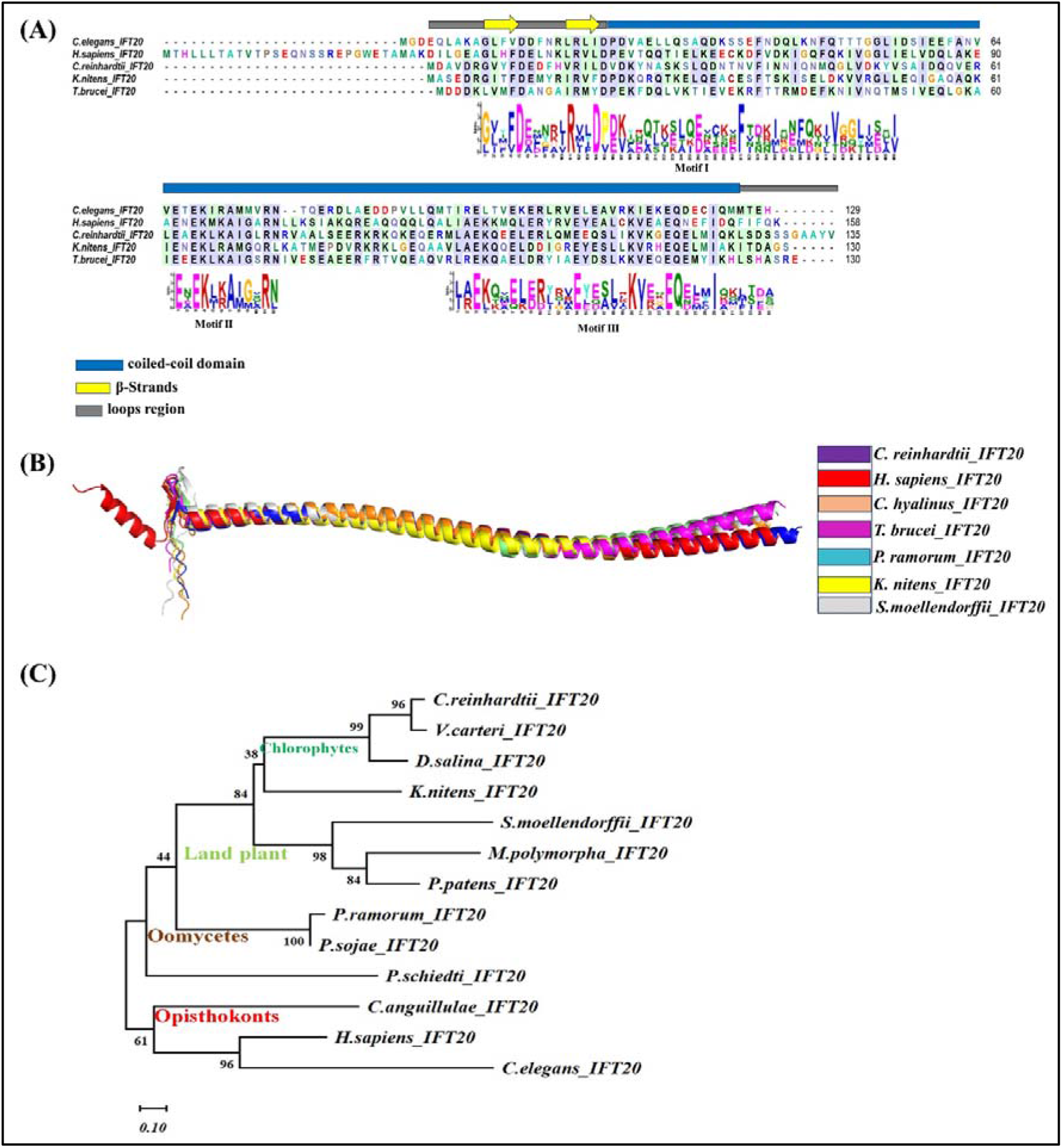
Sequence conservation and evolutionary relationships of IFT20 homologs. (A) Multiple sequence alignment of IFT20 protein sequences from diverse organisms. Conserved residues are highlighted in blue color shading. A schematic representation of the predicted secondary structural elements is shown above the alignment: α-helices (blue bars), β-strands (yellow arrows), and loop regions (grey). MEME (Multiple Expectation Maximization for Motif Elicitation) finds three discrete motif regions. (B) Structural superposition of IFT20 homologs colour coded by species. (C) Alignment of IFT20 protein sequences was carried out using ClustalW in MEGA11. The tree was analyzed using the Maximum likelihood method and the Jonas-Taylor-Thorton model with 1000 bootstrap replicates. Evolutionary analyses were conducted using MEGA 11 software. *(H. sapiens = Homo sapiens, T. brucei = Trypanosoma brucei, C. reinhardtii = Chlamydomonas reinhardtii, K. nitens = Klebsormidium nitens, C. elegans = Caenorhabditis elegans, P. patens = Physcomitrella patens, S. moellendorffii = Selaginella moellendorffii, C. hyalinus = Chytriomyces hyalinus*.

### 3.2 Cellular co-localization of IFT20-ChR1 complex in *C. reinhardtii*

For understanding IFT20-mediated ciliary trafficking of ChR1 in *C. reinhardtii*, we performed co-immunofluorescence detection using anti-CrIFT20 and anti-CrChR1-Ct antibody in the wild-type *C. reinhardtii* strain CC-125. The result shows that IFT20 (green) is prominently localized throughout the cilia. Co-immunostaining with ChR1 (red) demonstrated significant spatial overlap (yellow) with IFT20 in the merged fluorescence images, indicating that IFT20 and ChR1 are co-localized in the cilia (**Fig. 2A**). The immunofluorescence images show a distinct enrichment of ChR1 signal at the eyespot region, consistent with its known localization as a photoreceptor **(Fig. 2B)**. The eyespot appears as a concentrated, bright signal within the cell body. Notably, the IFT20 signal was also detected in the eyespot region, where the merged images revealed substantial overlap with ChR1, suggesting a potential spatial association at this site **(Fig. 2B)**. However, multi-cell images showed that eyespot co-localization was not consistently observed in all the cells, suggesting that this association is variable **(Supplementary Fig. S1)**. For the validation of the interaction between ChR1 and IFT20 proteins, co-immunoprecipitation was performed using anti-ChR1-Ct and anti-IFT20 antibodies in *C. reinhardtii* total cell lysate (CrTCL). ChR1 was found in the immunoprecipitant of IFT20 using anti-IFT20 antibody **(Fig. 2C)**. Similarly, in the reciprocal blot, IFT20 was detected in the ChR1-Ct immunoprecipitant using anti-ChR1-Ct antibody **(Fig. 2D)**. This shows that IFT20 facilitate the trafficking and localization of photoreceptors, such as ChR1, within the cilia of *C. reinhardtii*.

**Fig. 2.**
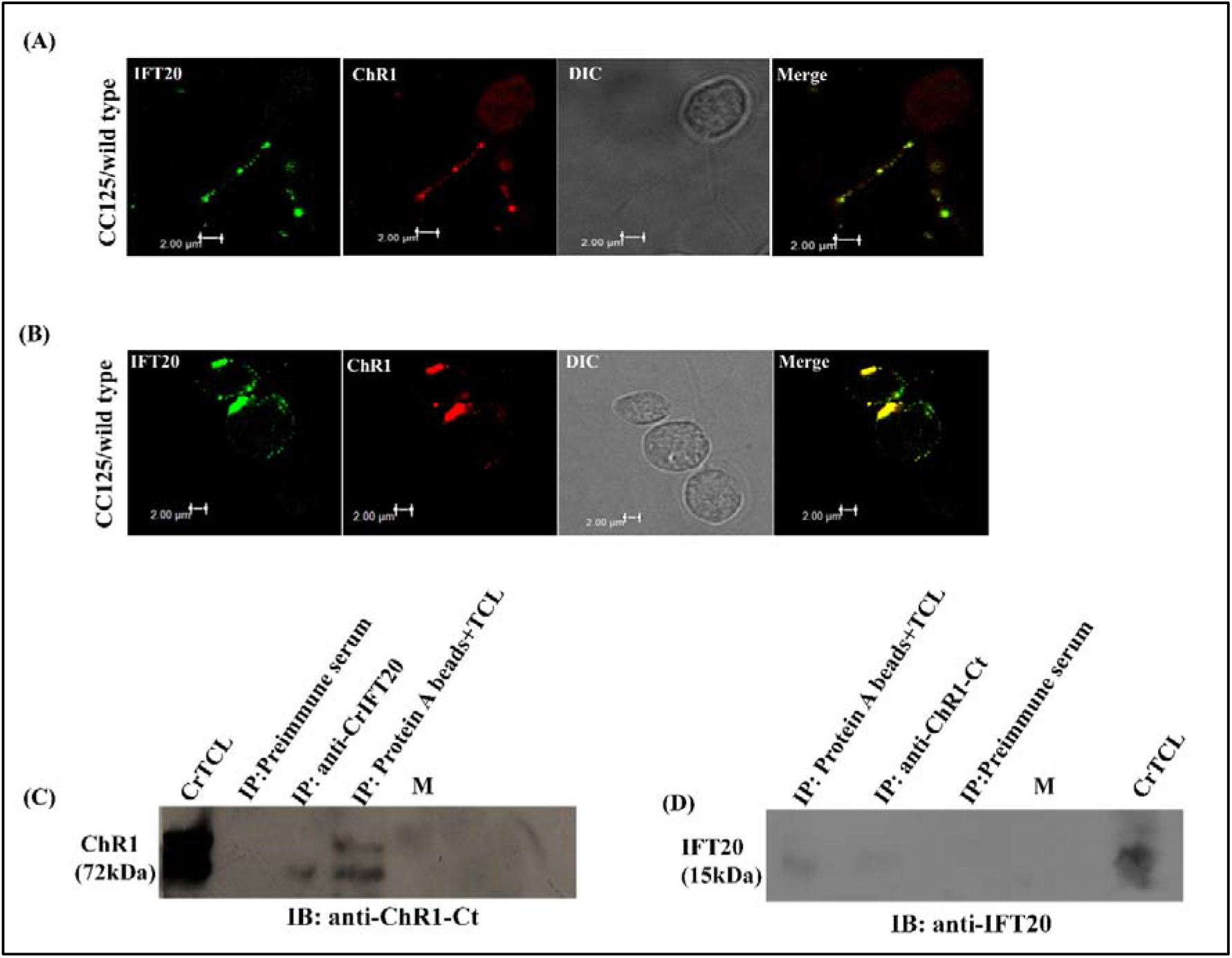
Co-localization of CrIFT20 with CrChR1 in *Chlamydomonas* wild type CC125. (A) Immunofluorescence staining of wild-type *Chlamydomonas reinhardtii* (CC-125). Images from left to right indicate fluorescence specific to CrIFT20 (green) and CrChR1 (red), Differential interference contrast (DIC), and merged images of the green and red channels showing colocalization (yellow). (B) Immunofluorescence of CrIFT20 (green), ChR1(red), merged (yellow) in *Chlamydomonas* in eyespots. Immunolocalization with anti-IFT20 antibody (1:250), anti-ChR1-Ct (1:250). Scale Bar = 2 μm. (C-D) Immunoprecipitation (IP) of CrIFT20 and CrChr1 using anti-IFT20 and ChR1-Ct antibody followed by immunoblotting (IB) with ChR1-Ct and IFT20 antibody. Protein bands of 72kDa represent ChR1 and 18kDa represent IFT20. IP with pre-immune serum served as negative control. (PIS = pre-immune serum).

### 3.3 BBS1 mutation results in altered ciliary distribution of IFT20-ChR1 complex

In the *bbs1* mutant, immunofluorescence analysis revealed altered localization patterns of IFT20 and ChR1. IFT20 (green) accumulated predominantly near the basal body region and along the cilia. In contrast, ChR1 (red) showed accumulation at the distal ends of the cilia rather than a uniform distribution as observed in wild-type cells. Differential interference contrast (DIC) imaging revealed normal cell morphology and full-length flagella, despite of *bbs1* mutation. The merged image shows co-localization of IFT20 and ChR1 at distal cilia, as indicated by the yellow signal **(Fig. 3)**. It was observed to be a multi-cell event **(Supplementary Fig. S2)**. Unlike wild-type cells, where ChR1 spreads evenly throughout the cilia, the *bbs1* mutant showed accumulation of ChR1 at the distal end, indicating that BBS1 protein plays a crucial role in the ciliary membrane protein trafficking along with IFT20.

**Fig. 3.**
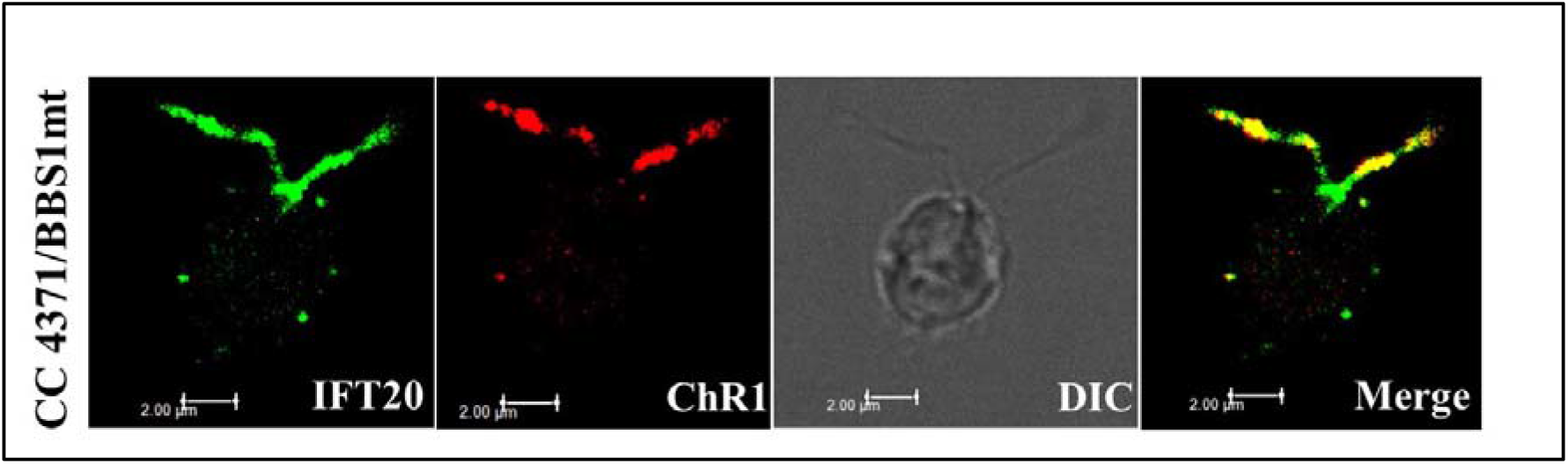
Mislocalization of the CrIFT20-CrChR1 complex in the CC-4371/BBS1 mutant **(A)** Immunofluorescence staining of CC4371/BBS1mt mutant. Left to right: showing CrIFT20 (green), CrChR1 (red), Differential interference contrast (DIC), merged (yellow). Scale bar = 2 μm.

### 3.4 Cellular co-localization of IFT20-Arf in cilia of *C. reinhardtii*

We identified the role of IFT20 in the trafficking of ChR1 in the cilia of *Chlamydomonas* wild type CC-125. To better understand IFT20-mediated ciliary trafficking, we have conducted a systems biology analysis in *C. reinhardtii*. Protein interactions network were predicted based on co-expression, gene fusion, gene neighborhood, and gene-occurrence. To identify potential protein complexes within the global network, we applied the Markov Clustering Algorithm (MCL). A total of five clusters were identified: Intraflagellar transport machinery, SNARE protein, Arf-like small GTPase, ER-to-Golgi anterograde transport, and photoreceptors. Hence, we also studied the role of IFT20 in the ciliary trafficking of the ancillary protein, Arf **(Fig. 4A)**. The co-immunofluorescence of IFT20 and Arf was performed in *C. reinhardtii* wild-type CC-125 cells. It was shown that IFT20 (green) co-localized with Arf in the *C. reinhardtii*’s cilia (yellow) **(Fig 4B)**. We also predicted the Alphafold multimer structure of the IFT20-Arf complex, however, the iPTM value for the predicted complex was found to be 0.55, suggesting a likely low confidence prediction (data not shown). Hence, the mechanism of the IFT20-mediated trafficking of Arf is not known and needs further experimental validation.

**Fig. 4.**
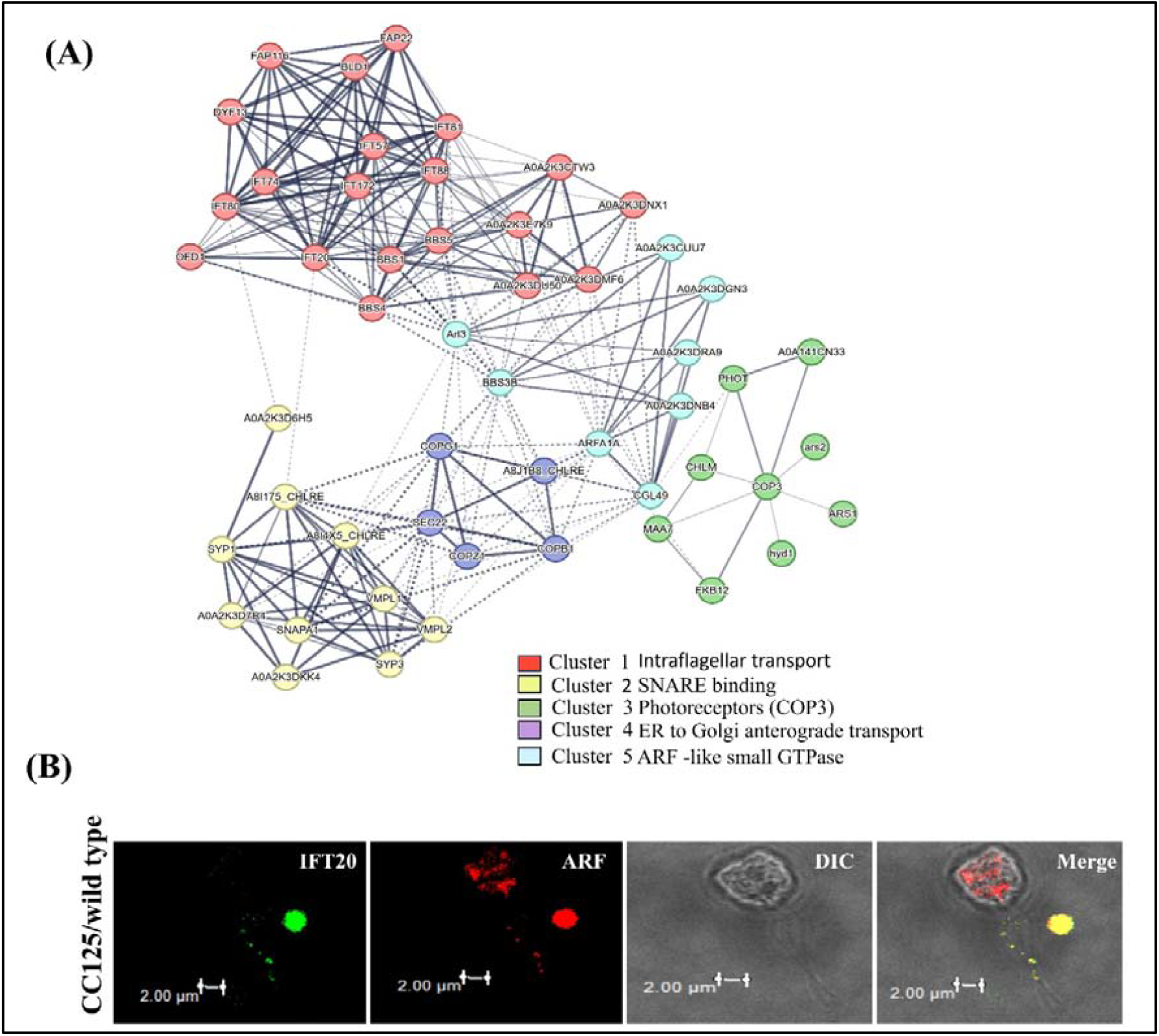
(A) Protein-protein interaction network predicted using STRING software in *C. reinhardtii*. IFT complex couples with the BBSome proteins, the ancillary ciliary protein, and photoreceptors to regulate ciliary membrane protein trafficking. Interactome analysis of IFT20 links components of the IFT complex, BBSome proteins, the vesicular trafficking system, and photoreceptors, underscoring its role in regulating cargo transport to cilia. Clusters are highlighted in colour to show variation in the interactome. (B) Cellular co-localization of CrIFT20 and Arf in *Chlamydomonas* wild type CC-125 cells. Immunofluorescence images showing the distribution of IFT20 (green) and Arf (red). Differential interference contrast (DIC) with merged channel shows IFT20 and Arf signal in cilia (yellow). Scale bar = 2 µm.

### 3.5 Biophysical characterization of recombinant IFT20

Recombinant CrIFT20 was expressed in *E. coli* using a heterologous system. The recombinant IFT20 was biophysically characterized using fluorescence spectroscopy and far-UV CD spectrophotometry. The intrinsic tyrosine fluorescence spectroscopy indicated a singular broad emission spectrum **(Fig. 5A)**. The spectrum of the purified IFT20 was also acquired following the increasing concentration of GTP (in 100 μM increments). A sequential quenching in fluorescence intensity was noted indicating the occurrence of GTP binding **(Fig. 5A)**. The far UV-CD data revealed that negative peaks at 208 nm and 222 nm for the recombinant IFT20 protein affirmed the presence of an alpha helical structure **(Fig. 5B)**. The change in far-UV CD spectra upon GTP addition aligned with the notion that the IFT20 protein may show a conformational change upon binding to GTP. To investigate possible interactions with nucleotides, a model of the complex between CrIFT20, GTP, and Mg^2^□ was generated using AlphaFold **(Fig. 5C-D)**. This model indicates a potential association with the iPTM value 0.71 **(Fig. 5C-D)**. Analysis of secondary structures across IFT20 homologs revealed a consistently high helical content of 76 %-93 % **(Supplementary Table S2)**. In contrast, the amount of β-strand and turn structures varied across lineages, suggesting that structural variability is confined to peripheral regions and not directly involved in IFT-B complex assembly as shown in PDB file no. 8RUY. Collectively, these findings suggest that IFT20 features a highly conserved helical core that is evolutionarily maintained to uphold its architectural and scaffolding roles within ciliary transport systems. It also shows modular variation in non-core segments that may contribute to lineage-specific adaptations in IFT20 protein dynamics.

**Fig. 5.**
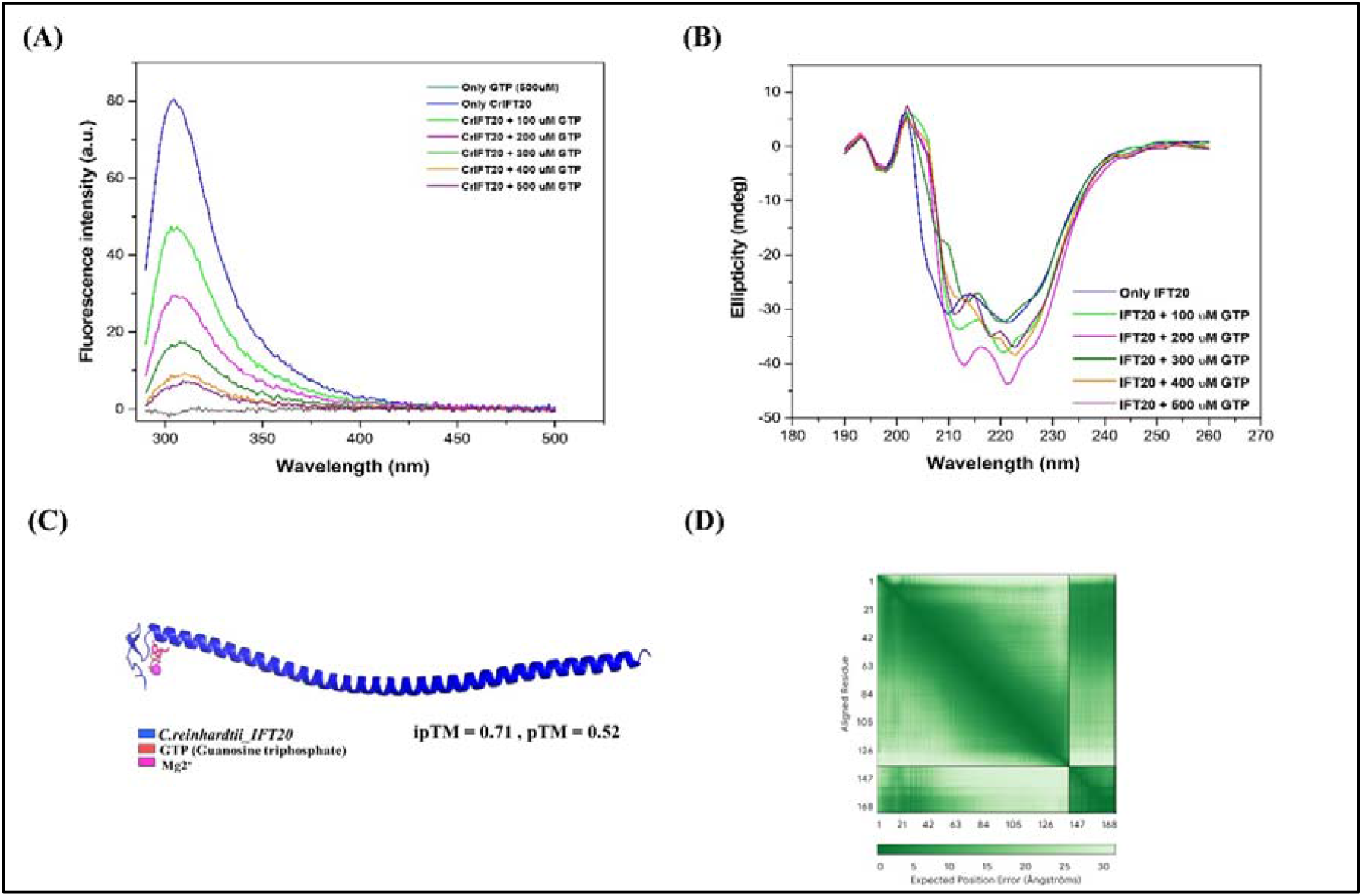
Structural comparison and spectroscopic characterization of recombinant CrIFT20 (A) The intrinsic fluorescence emission of recombinant CrIFT20 protein was recorded between 290-500 nm upon excitation at 275 nm. The single emission peak indicates the typical tyrosine fluorescence signature. (B) Circular dichroism (CD) spectrum of recombinant CrIFT20, with negative peaks characteristic of α-helical secondary structure. (GTP - Guanosine Triphosphate) (C-D) AlphaFold-multimer model of CrIFT20-GTP-Mg^2+^complex. Predicted Aligned Error (PAE) matrices indicate confidence in structural predictions for CrIFT20-GTP-Mg^2+^ complex.

## 4. Discussion

IFT20 plays a central role in ciliary trafficking, particularly in the delivery of membrane-associated proteins such as photoreceptors to the outer segment, as demonstrated in previous studies (Keady et al., 2011). Its dual localization to both the Golgi apparatus and the cilium further supports its function as a critical mediator linking early secretory pathways with intraflagellar transport (Follit et al., 2008). This spatial distribution positions IFT20 as a key coordinator of cargo sorting, delivery, and ciliary entry. Structural insights into IFT complexes provide an important framework for understanding IFT20 function (Lacey et al., 2023). Recent high-resolution models of IFT-A and IFT-B show that these complexes consist of elongated α-helical and coiled-coil subunits. These proteins create a flexible scaffold that allows the complexes to bind cargo and associate with the motors required for transport (Wachter et al., 2019, Lacey et al., 2024). Consistent with this architecture, our comparative sequence and phylogenetic analyses show that IFT20 contains a conserved C-terminal coiled-coil domain, likely essential for its integration into the IFT-B complex and for mediating protein-protein interactions. In contrast, the N-terminal region is less conserved and comprises β-strands connected by flexible loops, suggesting a role in species-specific interactions and regulatory adaptation (sequences for the analyses are mentioned in **Supplementary Table S3**. In addition to global structural features, specific sequence motifs appear to contribute to IFT20 function. The conserved YEFI motif has been implicated in mediating interactions with the WD40 domain of ATG16, linking IFT20 to autophagy-related pathways (Boukhalfa et al., 2021). Our analysis further identifies three discrete conserved motifs within the helical region, supporting the idea that these segments act as interaction interfaces within the IFT machinery. Together, these findings suggest that IFT20 combines a structurally conserved scaffold with adaptable regions. Further, the cellular characterization shows that IFT20 is evenly distributed throughout the cilia in wild-type CC-125 cells. The distribution of IFT20 along the cilia is consistent with its role in IFT trains, which mediate bidirectional cargo transport (Taschner and Lorentzen 2016). In wild-type CC-125 strain, co-immunoprecipitation (Co-IP) and co-immunofluorescence data show that IFT20 and ChR1 co-localize, suggesting that IFT20 is the vehicle for delivering photoreceptor ChR1 to the ciliary membrane. A particularly interesting observation is the localization of IFT20 at the eyespot. However, the fact that this co-localization of IFT20-ChR1 in the eyespot was not consistent across multiple cells implies that the IFT20-ChR1 association at the eyespot is a transient or regulated event, likely due to phototaxis. In the absence of the BBS1 protein, the IFT machinery still functions, as evidenced by IFT20 presence in the cilia. However, ChR1 fails to co-localize with IFT20 at the basal body, instead accumulating at the ciliary tips in the experimental setup condition. While IFT20 is sufficient to drive the anterograde (outward) movement of ChR1, the BBSome is clearly required to facilitate the proper sorting of the protein at the basal body. Without BBS1, the “exit” signal for ChR1 is lost, leading to the observed distal stagnation. By characterizing these differences between CC-125 and *bbs1* mutants, we show that ciliary health depends not just on the ability to move proteins in, but on the coordinated machinery required to move them around and out. The information regarding *C. reinhardtii* strains is mentioned in **Supplementary Table S4**. To better understand the cilia-trafficking context of these interactions, we mapped the protein-protein interaction network of IFT20 using STRING analysis. The resulting network reveals that IFT20 sits at a critical intersection of clusters of the intraflagellar transport machinery, the BBSome complex, and ancillary proteins (Arf-like small GTPase). Most notably, IFT20 shows a high degree of connectivity with the IFT complex and the BBSome. The co-localization of IFT20-Arf in cilia suggests the role of IFT20 in the trafficking of ancillary proteins. It is studied that Arf4 is localized to photoreceptor cells and plays a crucial role in the regulation of the protein trafficking to and within the cilium. These proteins manage the transport of essential cargo, such as rhodopsin, from the trans-Golgi network (TGN) to the photoreceptor outer segment (connecting cilium) (Deretic et al., 2005). Hence, mechanistic understanding of IFT20’s role in ancillary ciliary protein trafficking is further needed. Since Arf (small GTPase) co-localizes with IFT20 in cilia of CC-125 strain, we *in vitro* analyzed the probable interaction and binding of GTP with IFT20. The biochemical characterization showed that only the recombinant IFT20 protein has an α-helical structure. The far-UV CD data, along with the high helical content (76%-93%) predicted across IFT20 homologs, indicate an evolutionarily conserved core. This structural rigidity is likely essential for IFT20 to act as a stable “bridge” within the IFT-B complex, maintaining the scaffold’s integrity as it moves cargoes along the axoneme. The GTP-induced fluorescence quenching and the resulting shifts in the CD spectra suggest that IFT20 may bind to the GTP during probable IFT20-Arf interaction. Notably, CrIFT20 has no canonical GTPase switch region, however, it may bind to the GTP making it a non-canonical GTP-binding protein. However, the functional role of this interaction still remains unknown.

## 5. Conclusions

This study identifies *C. reinhardtii* IFT20 as a conserved adaptor that mediates ciliary trafficking of ChR1. The integrative analysis demonstrates that CrIFT20 possesses a highly conserved coiled-coil core, essential for its architectural role in IFT-B assembly and for persistent trafficking of membrane proteins within cilia. Comparative sequence, structural, and phylogenetic investigations reveal strong evolutionary conservation in the coiled-coiled domain required for interaction with key transport machinery. Cellular analyses further showed that IFT20 localizes within the cilia and co-localizes with the photoreceptor ChR1, as supported by co-immunoprecipitation. These observations identify IFT20 as a cargo-proximal module that facilitates the targeted delivery of receptors to the ciliary membrane.

## CRediT authorship contribution statement

**Suneel Kateriya:** Conceptualisation, Data curation and Methodology, Supervision, Validation, Funding acquisition, Writing-review and editing. **Alka Kumari**: Formal analysis, Investigation, Writing-original draft, Methodology. **Shilpa Mohanty:** Investigation, Writing-review and editing. **Snigdha Samanta:** Methodology

## Funding sources

We are thankful to ANRF/SERB, Government of India for granting EEQ/2023/000398 research project.

## Declaration of Competing Interest

The authors declare no conflict of interest.

## Acknowledgements

The research grant, EEQ/2023/000398from ANRF/SERB is duly acknowledged. Alka Kumari is the recipient of the Senior Research Fellowship from UGC (JRF), Govt. of India. The authors would like to thank Dr. Kumari Sushmita, Dr. Sunita Sharma and the lab members for their kind help. The SAITHI Foundation IIT Delhi and USIC, University of Delhi North Campus are acknowledged for providing instrumentation facilities.

## Data availability

The data will be available on request.

## Notes

### Competing Interest Statement

The authors have declared no competing interest.

### Summary of Updates

New experiments have been conducted to visualize the co-localization of IFT20 with the small GTPase, Arf protein in the cilia of Chlamydomonas reinhardtii. The manuscript has been further revised to make it more concise.

