## Supplementary Material for "Intraflagellar transport-20 mediates the ciliary membrane trafficking of channelrhodopsin in *Chlamydomonas reinhardtii*"

**Supplementary Figure S1**

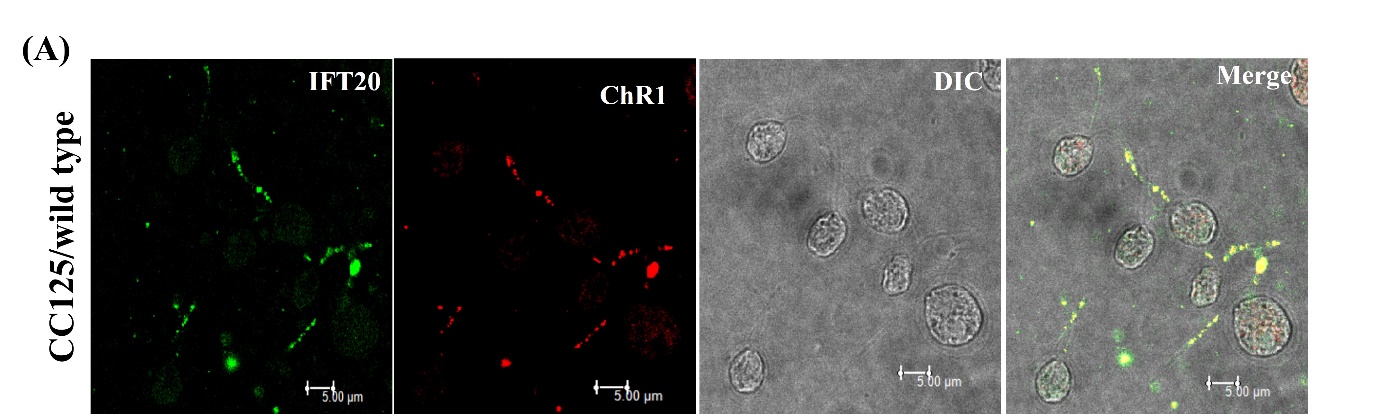

**Supplementary Fig. S1** Co-localization of CrIFT20 with CrChR1 in *Chlamydomonas* wild type CC-125. (A) Immunofluorescence staining of wild-type *Chlamydomonas reinhardtii* (CC-125) showing differential interference contrast (DIC) image with two intact flagella with the cell body, CrIFT20 immunostaining (green), CrChR1 immunostaining (red), and merged channels. CrIFT20 and ChR1 are both visible within the cilia, with significant spatial overlap in the merged image (yellow). Scale bar = 5 µm

**Supplementary Table S1** Predicted TM-score (pTM) values from pairwise structural alignment of IFT20 homologs

| **Organism** | **Predicted template modeling score (pTM)** |
| --- | --- |
| ***C.reinhardtii_IFT20*** | **0.53** |
| ***H.sapien_IFT20*** | **0.54** |
| ***K.nitens_IFT20*** | **0.52** |
| ***T.brucei _IFT20*** | **0.52** |
| ***P.ramorum_IFT20*** | **0.52** |
| ***S.moellendorffii_IFT20*** | **0.53** |
| ***C.hyalinus_IFT20*** | **0.53** |

**Supplementary Figure S2**

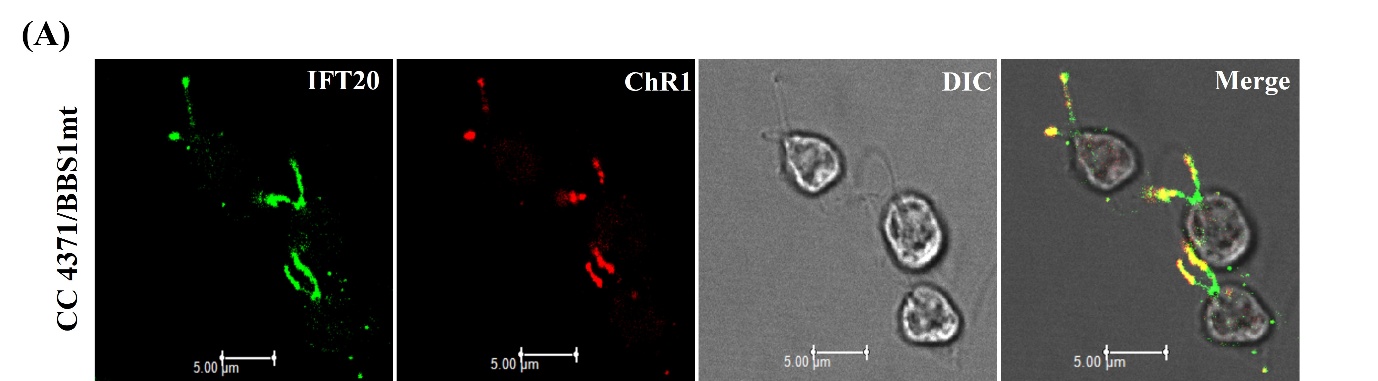

**Supplementary Fig. S2** Co-localization of CrIFT20 with CrChR1 in *Chlamydomonas* CC-4371 (*bbs1*-mt). Immunofluorescence staining showing differential interference contrast (DIC) image with two intact flagella with the cell body, CrIFT20 immunostaining (green), CrChR1 immunostaining (red), and merged channels (yellow). Scale bar = 5 µm

**Supplementary Table S2** Predicted secondary structure distribution of IFT20 homologs

|  | **Helix (%)** | **Sheet (%)** | **Turns (%)** |
| --- | --- | --- | --- |
| ***C.reinhardtiiI_FT20*** | **86.7** | **54.8** | **12.6** |
| ***V.carteri_IFT20*** | **85.2** | **57.8** | **12.6** |
| ***K.nitens_IFT20*** | **93.1** | **66.9** | **13.8** |
| ***P.patens_IFT20*** | **85.9** | **57.8** | **12.6** |
| ***P.sojae_IFT20*** | **76.7** | **61.7** | **17.3** |
| ***C.hyalinus_IFT20*** | **90.9** | **59.8** | **11.4** |
| ***T.brucei_IFT20*** | **86.2** | **33.1** | **14.6** |
| ***C.elengans_IFT20*** | **92.2** | **37.2** | **13.2** |
| ***H. sapiens_IFT20*** | **91.1** | **36.7** | **11.4** |

**Supplementary Table S3** List of sequences used in multiple sequence alignment, phylogenetic analysis, and structural superimposition.

| ***C.reinhardtiiI_FT20*** | MDAVDRGVYFDEDFHVRILDVDKYNASKSLQDNTNVFINNIQNMQGLVDKYVSAIDQQVERLEAEKLKAIGLRNRVAALSEERKRKQKEQERMLAEKQEELERLQMEEQSLIKVKGEQELMIQKLSDSSSGAAYV |
| --- | --- |
| ***V.carteri_IFT20*** | MDAADRGVYFDEDFHVRILDVDKYTASKSLQDNTNVFISNINNLQSLVEKYVSAIDQQVERLEAEKLKAIGLRNKVAALSEERKRKQKEQERMLGEKREELERLQMEEQSLIKVMGEQEILIQKLTDSSSGAAYV |
| ***D. salina_IFT20*** | MKFLDNTNIFITNVQQLTALVEKYLSQIDQQVERIEAEKLRAVGLRNRVAALEEERRRKHKEQERLIGEKQEELERLMTEEQSLMKVKQEQELLISKLSDSSSGAAFD |
| ***K.nitens_IFT20*** | MASEDRGITFDEMYRIRVFDPDKQRQTKELQEACESFTSKISELDKVVRGLLEQIGAQAQKIENEKLRAMGQRLKATMEPDVRKRKLGEQAAVLAEKQQELDDIGREYESLLKVRHEQELMIAKITDAGS |
| ***S. moellendorffii_IFT20*** | MALVSIDEESCVRLLPTDTYARCHSVENSCAAFRSKVSSFNKLVKELLGQVGERASKIENAKLCAIGTRNLVRDEMEKRPQELQVVKEGIDRRKEELERLRVEHLSLMAVKQEQEALLAKLSSASRKY |
| ***M. polymorpha_IFT20*** | MECVGLSARTAVGGGQLVQPPVPVLLDDASRLRVLHPDLHANSRVVEKTCNLFRSKLLQFHSLVKNLLAHVDEQSKKIEAAKSQAVGTRNMMSAEVETRTELLRDQRNIISDKQEHLERLNAELKSLFMVKQEQEVLIAQLSESSLPVQ |
| ***P.patens_IFT20*** | MAGDEAGSSAVVVDDNGRLRVLDLETNSHSRLVAKSCDVFQCKLQDFQNLVKQLLDQANEKAQQIEDAKLSAVGMRNMVSAEIETRPEKLRDQARLIADKQEQLERLNAEYESLLTVKQEQEALIARLSGSSLQV |
| ***P. ramorum,_IFT20*** | MSRGDNQLPSICFDDDCQVRVLDKDNITHTQELDQESNQFATKLEEFHEIVKGVLEVMEGQAKRIEREKLKAIGQRNRVDSEVENRIRQKQMLELQIKEKKTELERYNLQFQSLTKIADEQQVLMDKLSNNEA |
| ***P.sojae_IFT20*** | MSRGDNQMPSICFDDDYQVRVLDKDNITHTQELEQESNQFATKLEEFHEIVKGVLEVMEGQAKRIEREKLKAIGQRNRVDSEVENRNRQKQMLELLIKEKKTELERYNLQFQSLSKIADEQQALMDKLSNNEA |
| ***C.hyalinus_IFT20*** | MIRNSEKDSLGVTFDEFSKIRILPSDQFEASDSMKDQCKEFTQQIEDFNAIVQIFLEMLQAKAEQIEAEK  LKAIGLRNRVETELETRKAKRQQLQLLIKERHAELHRLKIQSDSLENVSLEQLRLIEQLSSK |
| ***C. anguillulae_IFT20*** | MISQQNSNTAAVSFDDLGRLRLVPAPLVDASDKLREECRDFSSKVTEFSTLVTDILNVVEAKAREIEKEKLKAIALRTAVEREPARRTAELNRISFLLRERQMQLDRISGLTSRRASRLMGQYETYAKMEQDQQAVIDQLTAK |
| ***T.brucei_IFT20*** | MDDDKLVMFDANGAIRMYDPEKFDQLVKTIEVEKRFTTRMDEFKNIVNQTMSIVEQLGKAIEEEKLKAIGSRNIVESEAEERFRTVQEAQVRLREKQAELDRYIAEYDSLKKVEQEQEMYIKHLSHASRE |
| ***C.elengans_IFT20*** | MGDEQLAKAGLFVDDFNRLRLIDPDVAELLQSAQDKSSEFNDQLKNFQTTTGGLIDSIEEFANVVETEKIRAMMVRNTQERDLAEDDPVLLQMTIRELTVEKERLRVELEAVRKIEKEQDECIQMMTEH |
| ***H. sapiens_IFT20*** | MTHLLLTATVTPSEQNSSREPGWETAMAKDILGEAGLHFDELNKLRVLDPEVTQQTIELKEECKDFVDKIGQFQKIVGGLIELVDQLAKEAENEKMKAIGARNLLKSIAKQREAQQQQLQALIAEKKMQLERYRVEYEALCKVEAEQNEFIDQFIFQK |

**Supplementary Table S4** List of *Chlamydomonas reinhardtii* strains used for the study.

| **Strain name** | **References** |
| --- | --- |
| **CC124(Wild-type)** | https://www.chlamycollection.org |
| **CC4371(BBS1/mt)** | (Lechtreck et al., 2009) |
